# ABC transporters alter plant-microbe-parasite interactions in the rhizosphere

**DOI:** 10.1101/526582

**Authors:** Deborah Cox, Steven Dyer, Ryan Weir, Xavier Cheseto, Matthew Sturrock, Danny Coyne, Baldwyn Torto, Aaron G. Maule, Johnathan J. Dalzell

**Author notes:** Joint first authors (Deborah Cox, Steven Dyer).

## Abstract

Plants are master regulators of rhizosphere ecology, secreting a complex mixture of compounds into the soil, collectively termed plant root exudate. Root exudate composition is highly dynamic and functional, mediating interactions between plants and a wide range of beneficial / harmful soil organisms. Exudate composition is under selective pressure to diversify in response to pathogen perception, whilst maintaining interactions with beneficial organisms. However, crop domestication has exerted significant and unintended changes to crop root exudate composition, and we know very little about genotype - phenotype linkages that pertain to root exudates and rhizosphere interactions. Better understanding could enable the modulation of root exudate composition for crop improvement by promoting positive, and impeding negative, interactions. Root expressed transporters modulate exudate composition and could be manipulated towards the rational design of beneficial root exudate profiles. Using Virus Induced Gene silencing (VIGS), we demonstrate that knockdown of two root-expressed ABC transporter genes in tomato cv. Moneymaker, *ABC-G33* and *ABC-C6*, alters the composition of semi-volatile compounds in collected root exudates. Root exudate chemotaxis assays demonstrate that knockdown of each transporter gene triggers the repulsion of economically relevant *Meloidogyne* and *Globodera* spp. plant parasitic nematodes, which are attracted to control treatment root exudates. Knockdown of *ABC-C6* inhibits egg hatching of *Meloidogyne* and *Globodera* spp., relative to controls. Knockdown of *ABC-G33* has no impact on egg hatching of *Meloidogyne* spp. but has a substantial inhibitory impact on egg hatching of *G. pallida. ABC-C6* knockdown has no impact on the attraction of the plant pathogen *Agrobacterium tumefaciens,* or the plant growth promoting *Bacillus subtilis,* relative to controls. Silencing *ABC-G33* induces a statistically significant reduction in attraction of *B. subtilis*, with no impact on attraction of *A. tumefaciens*. *ABC-C6* represents a promising target for breeding or biotechnology intervention strategies as gene knockdown (−64.9%) leads to the repulsion of economically important plant parasites and retains attraction of the beneficial rhizobacterium *B. subtilis.* This study exposes the link between ABC transporters, root exudate composition, and *ex planta* interactions with agriculturally and economically relevant rhizosphere organisms, paving the way for an entirely new approach to rhizosphere engineering and crop protection.

## Introduction

Plants secrete a complex mixture of water soluble and volatile organic compounds (VOCs) into the soil, collectively termed plant root exudate. Root exudates can enhance the recruitment of beneficial microbes (Kim et al., 2016; Li et al., 2016; Allard-Massicotte et al., 2016; Yuan et al., 2015; Badri et al., 2009), mobilise nutrients (Waters et al., 2018), sequester toxic compounds in the soil (De Andrade et al. 2011), and communicate with other plants, and animals (Hiltpold et al. 2010; Bertin et al., 2003). Root exudate composition is dynamic, and can be modulated as a factor of development (Chaparro et al. 2013; Byrne et al., 2001), environment (Giles et al., 2017), physiological state (Badri et al., 2009), and displays marked diversity between species (Zwetsloot et al., 2018; Bowsher et al., 2016; Cieslinski et al., 1997; Fletcher and Hegde, 1995) and cultivars (Mohemed et al., 2018; Kihika et al., 2017; Monchgesang et al., 2017 & 2016; Micallef et al., 2009). It is estimated that between 5% and 21% of all photosynthetically-assimilated carbon is released as root exudate (Jones et al. 2009; Marschner, 1995).

The process of crop domestication has focused on a small number of desirable traits, relating to plant stature, yield and disease resistance (Meyer et al., 2012) often at the expense of other traits. For example, there is evidence that the domestication process has exerted a significant and unintended impact on root exudate composition, and rhizosphere microbe interactions (Iannucci et al., 2017; Bulgarelli et al., 2015). A general lack of understanding and mechanistic insight represents a major impediment to the exploitation of root exudates for crop improvement. However, it is clear that root exudate composition is an adaptive trait, which can be manipulated. For example, much progress has been made in understanding the interaction between root exudates, and parasitic *Striga* spp. over recent years. This insight has underpinned efforts to alter exudate strigolactone content, a known germination stimulant and attractant of these economically important and globally distributed parasitic plants (Mohemed et al., 2018; Xu et al., 2018; Gobena et al., 2017; Mohemed et al., 2016; Jamil et al., 2012). Comparatively, less progress has been made in understanding the analogous interaction between root exudate and plant parasitic nematodes (PPNs), in part due to the increased complexity of nematode biology. Nonetheless, recent years have seen renewed interest in this field of research (Čepulytė et al., 2018; Hoysted et al., 2018; Murungi et al., 2018; Kirwa et al., 2018; Kihika et al., 2017; Warnock et al., 2016).

It is estimated that PPNs reduce crop yields by 12.3%, equating to an estimated $US 80 - 157 billion in losses each year (Coyne et al., 2018; Jones et al., 2013; Nicol et al., 2011). PPNs respond transcriptionally, physiologically and behaviourally to plant root exudates, using exudates to trigger egg hatching, and to facilitate host-finding (Cepulyte et al., 2018; Palomares-Rius et al., 2016; Warnock et al., 2016; Yang et al. 2016; Zasada et al., 2016; Duarte et al., 2015; Teillet et al., 2013). Understanding the molecular, chemical and physiological mechanisms underpinning both root exudation and PPN interactions could facilitate the development of aggressive new *ex planta* control strategies for sustainable intensification of global agriculture, through breeding, rhizosphere engineering and / or biotechnology (Ahkami et al., 2017; Warnock et al., 2017; Dessaux et al., 2016; Devine and Jones, 2001). The identification of key parasite attractants and repellents could also facilitate the development of new push-pull strategies.

The rhizosphere microbiome is also a major contributing factor to crop health (Sasse et al., 2018; Berendsen et al., 2012), and phenotype (Hubbard et al., 2018; Lu et al., 2018). Microbial chemotaxis to plant root exudates is an important factor in the competition for chemical resources in the rhizosphere, and colonisation of plant roots (Allard-Massicotte et al., 2016; de Weert et al., 2002). As such, alteration of root exudate composition could impact on a wide range of interactions. Exploitation of root exudates for improved crop health offers intriguing potential, but requires a detailed study of the link between crop genotype and highly complex, multi-species interactions.

Considerable interest has developed around the manipulation of membrane transporters for crop improvement (Lane et al., 2016; Schroeder et al., 2013), and ABC transporters have been implicated directly in modifying root exudate composition (Badri et al., 2008, 2009). ABC transporters represent one of the single largest gene families in plants, which regulate the sequestration and mobilisation of a vast array of chemistry linked to diverse metabolic, physiological and morphological functions (Adebesin et al., 2017; Hwang et al., 2016; Martinoia et al., 2012; Yazaki et al., 2006; Liu et al., 2001). The ABC transporter gene complement of tomato is numbered at 154, with a considerable proportion expressed in root tissue (Ofori et al., 2018). Here we have employed an improved Virus Induced Gene Silencing (VIGS) method to reveal a functional link between two ABC transporter genes, root exudate composition, and rhizosphere interactions with economically important microbes and parasites.

## Results

### Knockdown of tomato ABC transporter genes by VIGS

VIGS was used to co-silence two genes of interest (*ABC-C6* or *ABC-G33*), alongside a visual reporter gene, Phytoene DeSaturase (*PDS*). Knockdown of *PDS* triggers a mild leaf bleaching phenotype (Winzer et al., 2012). Co-silencing was necessary to identify responsive plants for exudate collection and downstream bioassays; not all plants will trigger a viable RNAi response to VIGS challenge. We observed that plant growth rate was reduced by over 75% relative to control treatments when using the traditional blunt syringe inoculation method. We therefore sought to develop a less damaging approach to inoculation of *A. tumefaciens* (containing the VIGS plasmids). We discovered that *A. tumefaciens* cultures could efficiently invade leaf cells when applied topically to tomato seedling cotyledons with Silwett L-77, which is frequently used to aid *A. tumefaciens* invasion during floral dip transformation protocols (Clough & Bent, 1998). Following topical application of *A. tumefaciens* (containing the VIGS plasmids), we did not observe any reduction in plant growth rate, and bleaching phenotypes typically began to develop within 11 days and peaked at around 21 days post inoculation. Bleaching phenotypes following blunt syringe application began to emerge around 15 days post inoculation, and similarly peaked around 21 days post inoculation. Furthermore, the frequency of plants demonstrating the mild photobleaching phenotype associated with *PDS* knockdown was between 80% and 100% following topical application. The blunt syringe leaf infiltration method resulted in around 75% of plants demonstrating the co-silenced bleaching phenotype.

Both inoculation methods triggered robust and specific gene knockdown by week three post inoculation (Figure 1). Plants were sampled three weeks post inoculation for further experiments. Transcript abundance of *ABC-C6* and *ABC-G33* was reduced by 63.3% ± 11.6% (P<0.01**) and 66.3% ± 8.6% (P<0.001***) using blunt syringe inoculation. Topical application resulted in transcript knockdown of 64.9% ± 9.4% (P<0.01**) and 46.4% ± 5.8% (P<0.001***) for *ABC-C6* and *ABC-G33* respectively. Due to the improved performance of topical inoculation in terms of plant growth rate, we adopted this method for all subsequent experiments. Only plants that displayed co-silenced bleaching phenotypes were taken for further analysis across all experiments.

**Figure 1.**
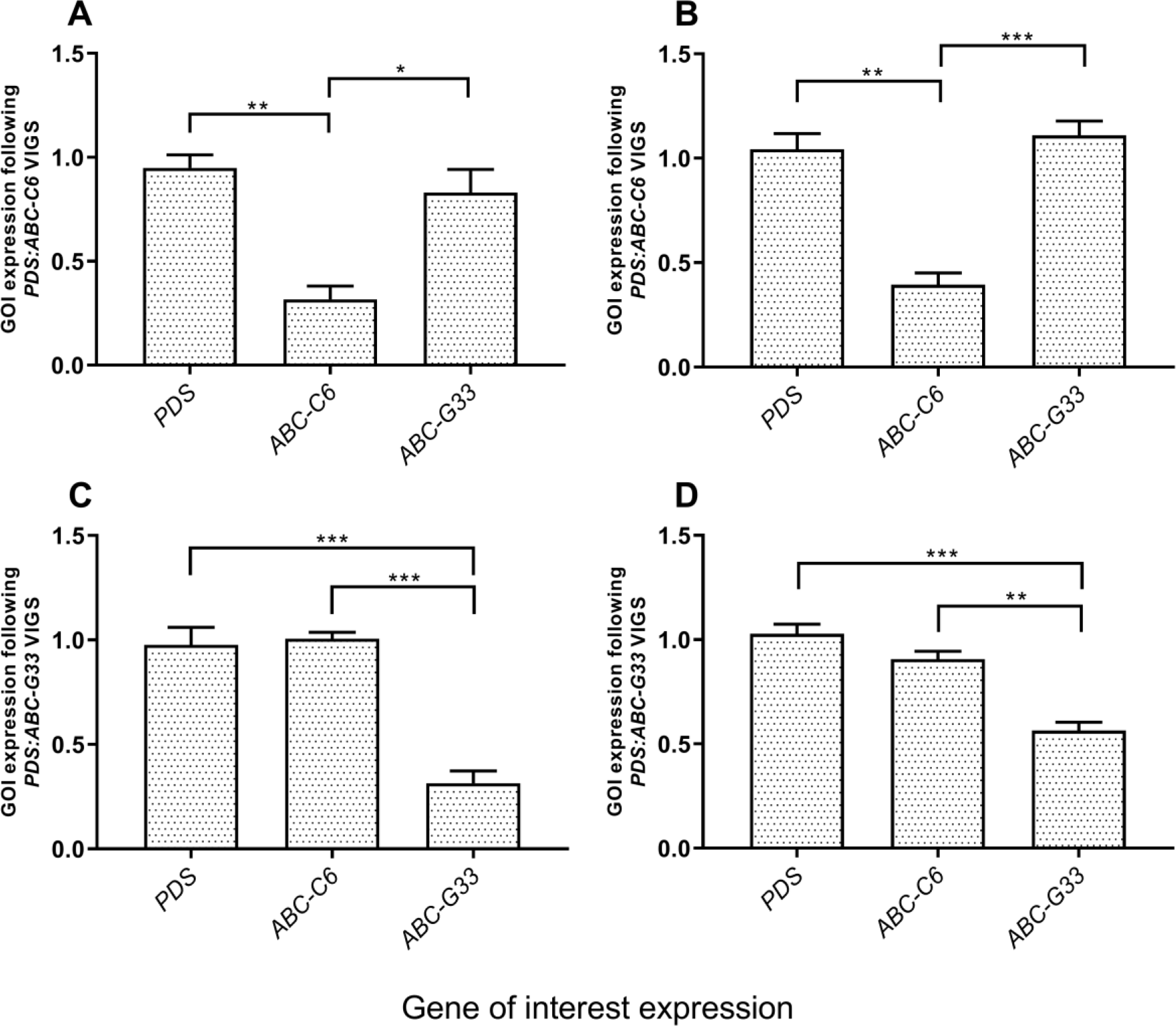
VIGS triggers target-specific knockdown of root-expressed ABC transporter genes in tomato. (A) The mean ratio of *ABC-C6* abundance relative to the endogenous control gene, elongation factor 1 subunit alpha (*EF-α*), following blunt syringe inoculation of *A. tumefaciens* and pTRV plasmids. (B) The mean ratio of *ABC-C6* abundance relative to the endogenous control gene following topical application of *A. tumefaciens* and pTRV plasmids. (C) The mean ratio of *ABC-G33* abundance relative to the endogenous control gene following blunt syringe inoculation of *A. tumefaciens* and pTRV plasmids. (D) The mean ratio of *ABC-G33* abundance relative to the endogenous control gene following topical application of *A. tumefaciens* and pTRV plasmids. Data represent three biological replicates, with each replicate consisting of three plants each; error bars represent SEM. One-way ANOVA and Tukey’s HSD tests were used to assess statistical significance between groups (P < 0.05*, P< 0.01**, P< 0.001***).

### Knockdown of ABC transporter genes modulate exudate composition and parasite behaviour

Root exudate was collected from VIGS treatment groups at three weeks post inoculation. Behavioural responses of two root knot nematodes, *Meloiodogyne incognita* and *Meloidogyne javanica,* alongside the potato cyst nematode, *Globodera pallida* were assayed across experimental exudates.

### Hatching

The hatching response of each species was measured as the percentage of emerging infective second stage juveniles (J2s) over time, for which the area under the curve was calculated for comparison of experimental groups (Figure 2C-E). Measuring the area under the curve allows for a more robust assessment of hatching phenotypes over time, as it is proportional to both the rate of hatching (gradient) and also the final hatch percentage. Knockdown of *ABC-C6* and *ABC-G33* triggered a reduction in final hatch at day 21 of 11.6% ± 3.4 (P<0.05*) and 36.2% ± 6.0 (P<0.001***) respectively for *G. pallida*. Area under the cumulative percentage hatch (AUCPH) was also reduced following knockdown of *ABC-C6* and *ABC-G33* by 174.4 ± 22.04% days (P<0.001***) and 420.1 ± 23.29% days (P<0.0001****), respectively. Emergence of *Meloidogyne* spp. J2s was assayed by measuring the ratio of hatched : unhatched J2s in each treatment over time, and converting to a percentage. The AUCPH was reduced for *M. incognita* following knockdown of *ABC-C6,* by 33.8 ± 7.3% days (P<0.01**). Knockdown of *ABC-G33* caused a hatching reduction of 16.9 ± 7.1% days (P>0.05, ns). Knockdown of *ABC-C6* also triggered a reduction in hatching of *M. javanica* of 112.4 ± 33.3% days (P<0.05*), whereas knockdown of *ABC-G33* led to a reduction of 59.7 ± 30.8% days (P>0.05, ns).

**Fig. 2.**
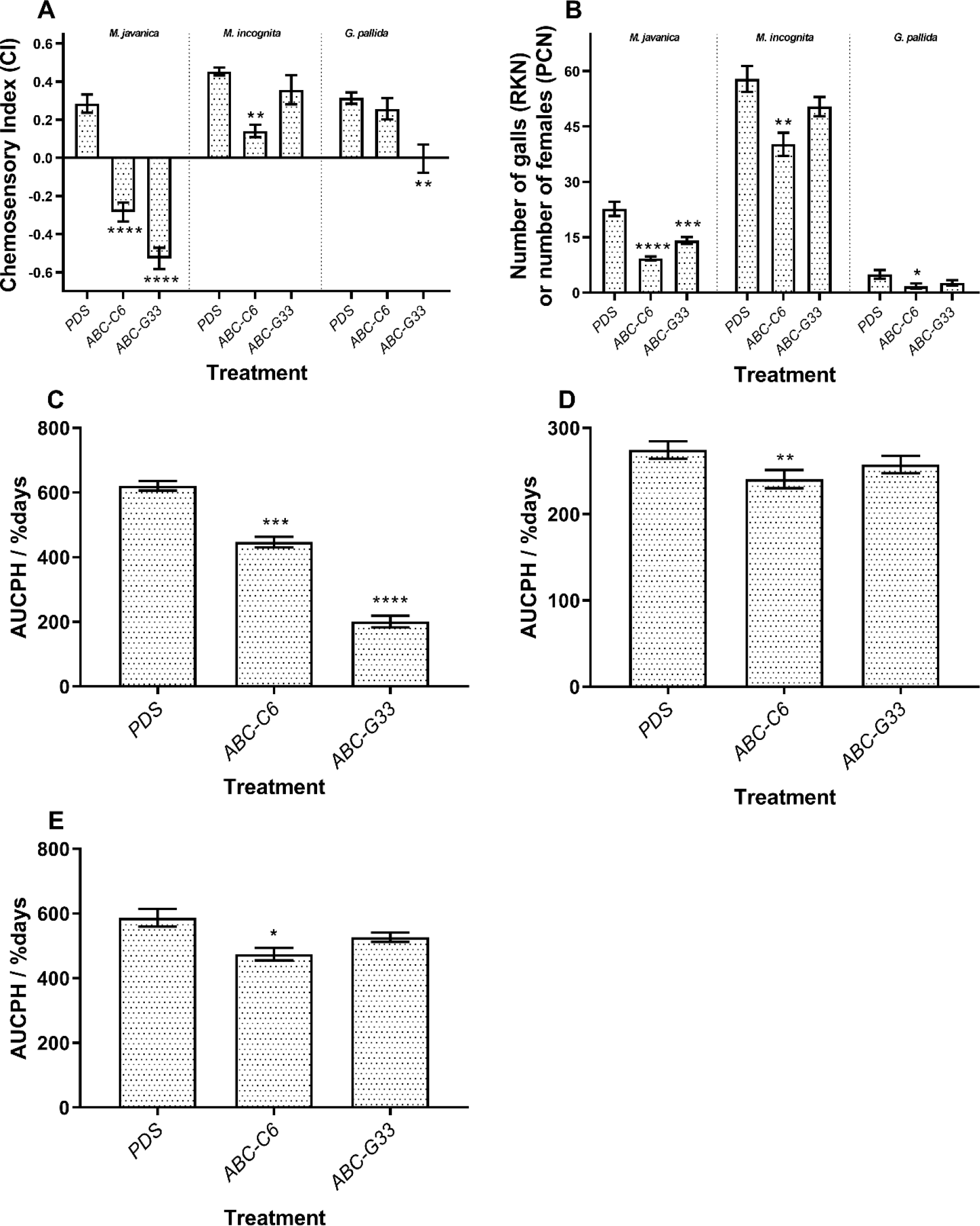
ABC transporter gene knockdown modulates PPN hatching, attraction and invasion. (A) Chemosensory challenge of *Meloidogyne* and *Globodera* spp. A positive chemosensory index (CI) equates to attraction towards tested exudates, and a negative CI infers repulsion from the exudates. Data represent five biological replicates for each nematode species in each treatment group. (B) The number of galls formed (*Meloidogyne* spp.) or developing females found on the root surface (*G. pallida*) six weeks post inoculation on VIGS plants (n=10 plants per species). (C) *G. pallida* J2 hatching was measured as the % of total eggs hatched at two-day intervals over three weeks at 17°C in darkness when incubated in experimental root exudates (n=4). The area under the curve from the cumulative percentage hatch (AUCPH) was estimated by trapezoidal integration, as described by Campbell & Madden (1990). (D) *M. incognita* J2 hatching was calculated as the observed ratio of hatched to unhatched J2 in a suspension of experimental exudate samples. The area under the curve (AUC) for these ratios was calculated using trapezoidal integration. (E) Hatching of *M. javanica* was calculated as above. Error bars represent SEM. Asterisks indicate statistical significance relative to controls following one-way ANOVA and Tukey’s HSD tests: P<0.05*; P< 0.01**; P< 0.001***; P< 0.0001****.

### Attraction

Knockdown of *ABC-C6* modified the chemosensory responses of *M. incognita* and *M. javanica* to collected exudates (Figure 2A). Experimentally manipulated exudates were less attractive to *M. incognita* J2s, whereas *M. javanica* J2s were repelled by the same exudates. Following knockdown of *ABC-C6*, chemosensory indices (CI) were reduced by 0.31 ± 0.07 (P<0.01**) and 0.57 ± 0.07 (P<0.0001****) for *M. incognita* and *M. javanica* respectively, relative to PDS control exudates. Following knockdown of *ABC-G33*, *M. javanica* was consistently repelled by these exudates with a CI score reduction of 0.76 ± 0.11(P<0.0001****), whereas *M. incognita* retained attraction. The CI score for *M. incognita* to *ABC-G33* exudates was reduced by just 0.1 ± 0.01 (P>0.05, ns). *G. pallida* J2s displayed reduced attraction to exudates from *ABC-G33* knockdown plants, by 0.32 ± 0.08 (P<0.01**), whereas for *ABC-C6* exudates, the CI was reduced by only 0.06 ± 0.06 (P>0.05, ns).

### Plant infection

Knockdown of ABC transporter genes also reduced the number of galls produced following infection of VIGS plants with *Meloidogyne* spp. (Figure 2b); knockdown of *ABC-C6* and *ABC-G33* caused a reduction of 17.7 ± 4.4 galls (P<0.0001****) and 7.5 ± 4.3 galls (P>0.05, ns) respectively, following infection with *M. incognita.* The number of galls produced following infection by *M. javanica* was reduced by 13.4 ± 1.8 (P<0.0001****) and 8.6 ± 1.8 (P=0.0005***) when *ABC-C6* and *ABC-G33* knockdown plants were challenged, respectively. The number of *G. pallida* females was reduced by 3.2 ± 1.2 following knockdown of *ABC-C6* (P<0.05*), whereas knockdown of *ABC-G33* caused a decrease of 2.3 ± 1.1 cysts per plant (P>0.05, ns). *G. pallida* retained attraction to exudates collected following *ABC-C6* knockdown, however cyst counts were significantly reduced following infection of VIGS plants, indicating that *in planta* consequences of transporter dysregulation can be distinct from *ex planta* implications (Figure 2c).

### Metabolomic characterisation of exudates following ABC transporter knockdown

Collected exudates were assessed by coupled gas chromatography-mass spectrometry (GC-MS) to identify changes in exudate composition. Several compounds were quantitatively altered in exudates collected following ABC gene knockdown, relative to *PDS* knockdown controls (Fig 3). Knockdown of *ABC-C6* resulted in reduced abundance of 2-methyloctacosane (P<0.001***), and increased abundance of nonadecane (P<0.01**) and tetradecanoic acid (P<0.05*), relative to control treatment (*PDS* knockdown). Contrastingly, knockdown of *ABC-G33* triggered elevated abundance of eicosane (P<0.001***), 9-O-pivaloyl-N-acetylcolchinol (P<0.001***), heptadecane (P<0.05*) and octadecanoic acid (P<0.05*).

**Figure 3.**
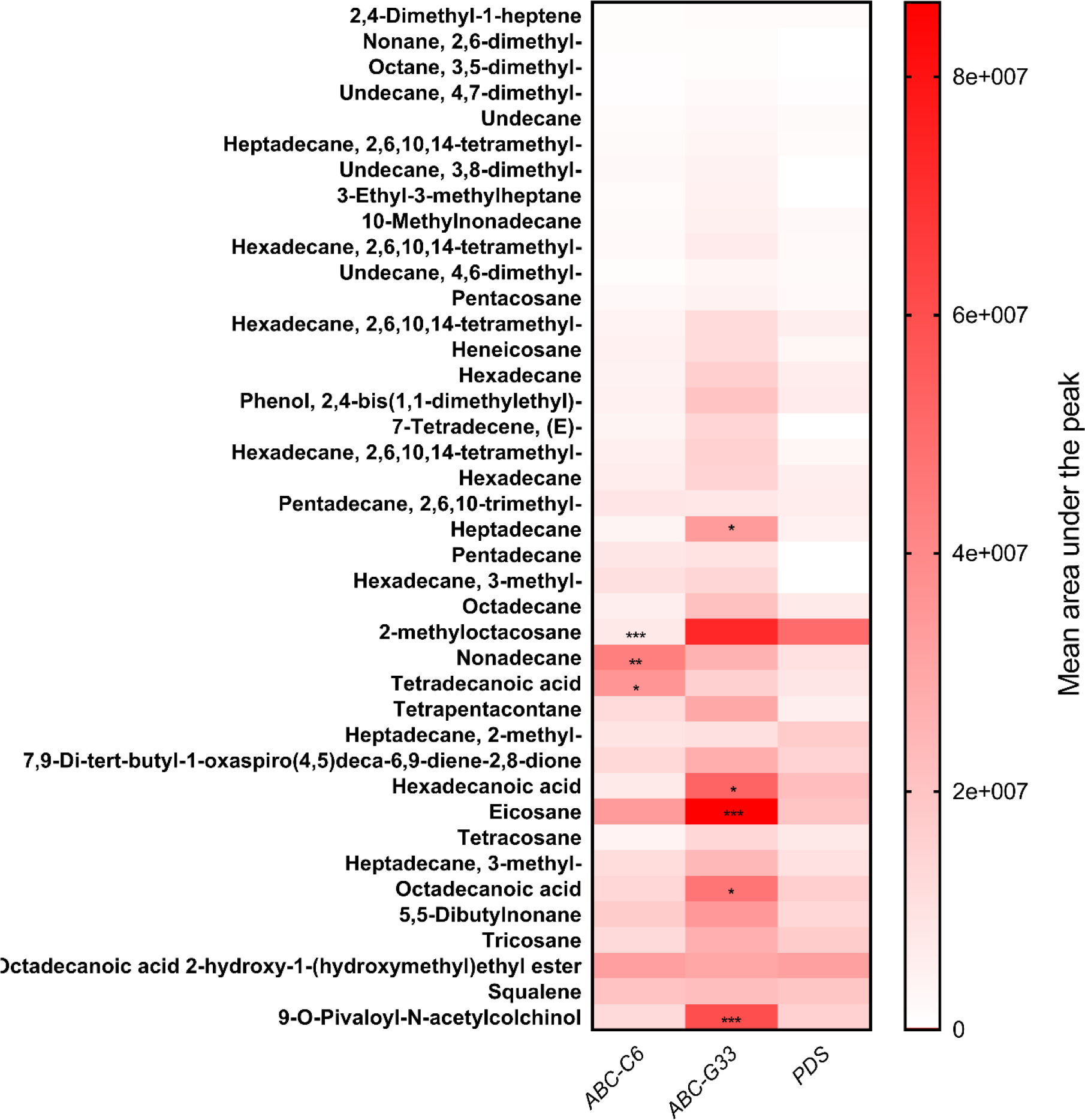
Heatmap showing differences in the relative abundance of identified compounds across experimental exudates. The mean composition of 10 biological replicates (three plants per replicate) is plotted for each experimental group post-VIGS, and has been assessed by two-way ANOVA, and Tukey’s multiple comparison test. Statistical significance is indicated relative to the *PDS* knockdown control, P<0.05*, P<0.01**, P<0.001***.

### Parasite behavioural responses to selected differentially exuded compounds

PPN species were assayed for responsiveness to selected compounds that were differentially exuded following knockdown of either ABC transporter gene. Tetradecanoic acid, hexadecanoic acid, octadecanoic acid and pentadecane were solubilised in 100% dimethyl sulfoxide (DMSO) (1 mM stocks). Each solubilised compound was independently inoculated into control exudates (*PDS* knockdown) to a final experimental concentration of 1 μM 0.1% DMSO. Inoculated control exudates were then used for egg hatching and chemotaxis assays (Figure 4).

**Figure 4.**
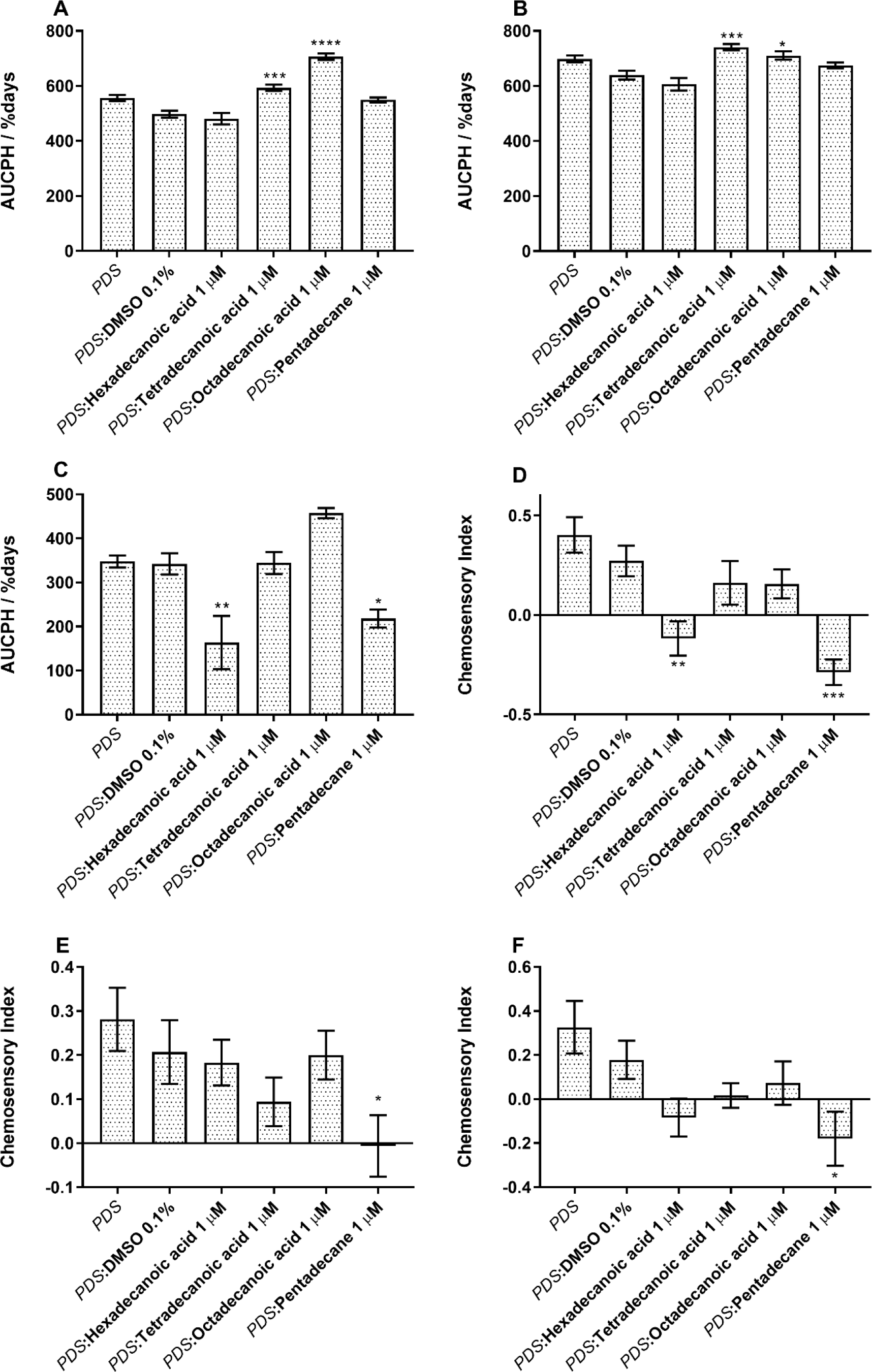
PPN responses to selected exudate compounds. Hatch responses following inoculation of control exudate with selected compounds for: (A) *M. incognita*; (B) *M. javanica*; (C) *G. pallida*. The area under the curve of percentage hatch (AUCPH) was estimated by trapezoidal integration and compared by one-way ANOVA and Dunnett’s multiple comparisons test and asterisks indicate statistical significance in hatching relative to the PDS:DMSO control. Chemotaxis responses to selected compounds inoculated into control plant root exudate (*PDS* knockdown treatment) for: (D) *M. incognita*; (E) *M. javanica*; (F) *G. pallida*. Data represent the mean of five biological replicates of root exudate, for which eight replicate assays are performed for each nematode species. Asterisks indicate statistical significance in chemosensory index relative to the PDS:DMSO control following one-way ANOVA and Tukey’s HSD tests. P< 0.01**; P< 0.0001****, error bars represent SEM. P< 0.001***; P< 0.0001****, error bars represent SEM.

Egg hatching of *Meloidogyne* spp. was enhanced when tetradecanoic acid or octadecanoic acid were inoculated into *PDS* knockdown exudates. The AUCPH for *M. incognita* increased by 95 ± 16.82% days (P<0.001***) and 208.1 ± 17.22% days (P<0.0001****), respectively (Figure 4A). For *M. javanica* the AUCPH increased by 101.7 ± 19.52 (P<0.001***) and 71.2 ± 21.76 (P<0.05*), respectively (Figure 4B). However, the addition of hexadecanoic acid and pentadecane significantly inhibited egg hatching in *G. pallida* by 178.7 ± 65.2% days (P<0.05*) and 124.2 ± 31.8% days (P<0.01**), respectively (Figure 4c).

The addition of 1 μM pentadecane to control root exudates reduced the CI of *M. incognita* by 0.56 ± 0.10 (P<0.01**). Likewise, the addition of 1 μM hexadecanoic acid reduced the CI of *M. incognita* by 0.39 ± 0.12 (P<0.001***). 1 μM tetradecanoic acid, or octadecanoic acid had no statistically significant impact on *M. incognita* attraction to control root exudates. For *M. javanica*, a 0.21 ± 0.1 (P<0.05*) decrease in CI was observed upon addition of 1 μM pentadecane; no statistically significant differences were observed following addition of the other compounds. The CI of *G. pallida* to root exudates was reduced by 0.36 ± 0.15 (P<0.05*) with the addition of 1 μM pentadecane.

### Knockdown of ABC transporter genes selectively modulates microbial chemotaxis

The attraction of *B. subtilis* and *A. tumefaciens* to root exudates was assessed following ABC gene knockdown. Both species were significantly more attracted to the positive control, 1 mM malic acid, relative to the negative ddH_2_O control. Similarly, root exudates collected from the *PDS* knockdown plants were significantly more attractive to both *B. subtilis* (P<0.0001****) and *A. tumefaciens* (P<0.001***), than ddH_2_O (Figure 5). The attraction of *B. subtilis* to exudates collected following knockdown of *ABC-C6* was statistically unchanged relative to the control treatment. However, knockdown of *ABC-G33* triggered a reduced attraction, and a mean reduction of 98.9 colony forming units (CFUs) relative to exudates from *PDS* knockdown plants (P<0.01**). Knockdown of *ABC-C6* and *ABC-G33* had no impact on the attraction of *A. tumefaciens* to collected exudates.

**Figure 5.**
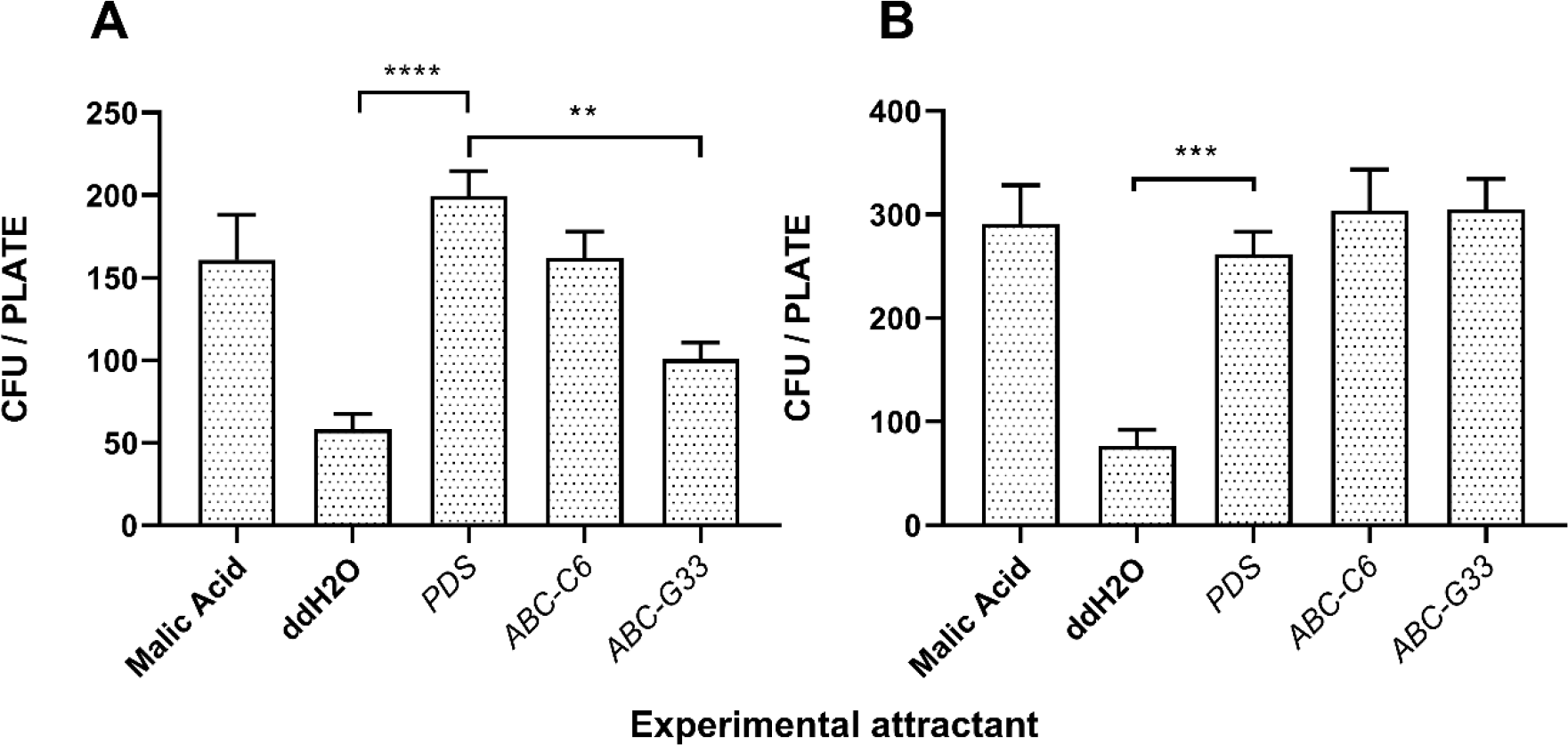
ABC transporter gene knockdown modulates the attraction of *B. subtilis,* but not *A. tumefaciens*. (A) Chemosensory response of *B. subtilis* (168) to root exudates collected following gene knockdown. (B) Chemosensory response of *A. tumefaciens* (AGL-1) to root exudates collected following gene knockdown. Data represent ten biological replicates for each species. P<0.01**; P< 0.001***; P<0.0001****, error bars represent SEM.

## Discussion

The predominantly sessile life-style of a plant necessitates substantial molecular and biochemical plasticity to coordinate responses to environmental conditions, and to interact contextually with floral and faunal communities. It is clear that plant genotype influences exudate composition, and organismal interactions (Mohemed et al., 2018; Iannucci et al., 2017; Kihika et al., 2017; Monchgesang et al., 2017 & 2016; Bulgarelli et al., 2015; Micallef et al., 2009). However, our understanding of the basic biology underpinning these interactions is limited.

ABC transporters modulate root exudate composition, and rhizosphere microbe interactions (Badri et al., 2008, 2009, reviewed in Sasse et al,. 2018). In this study, we used VIGS to investigate the role of two ABC transporter genes, *ABC-C6* and *ABC-G33,* in modulating tomato root exudate composition, and interactions with three important PPN species. Specifically, we demonstrate that knockdown of *ABC-C6* and *ABC-G33* transporter genes quantitatively alters tomato root exudate composition and inhibits PPN hatching and attraction behaviours to varying degrees. We assessed the involvement of individual compounds in mediating interactions with the PPN species by inoculating selected, differentially exuded compounds into control exudates. By assessing hatching and chemotaxis responses to these experimentally manipulated exudates, we demonstrate that hexadecanoic acid and pentadecane are inhibitors of *G. pallida* host-finding and hatching; pentadecane is an inhibitor of *Meloidogyne* spp. host-finding. Tetradecanoic acid and octadecanoic acid had no impact on host-finding of either species but did enhance egg hatching rates of both *Meloidogyne* spp. These results suggest that hydrocarbons and fatty acids in root exudate both mediate PPN host finding. However, our findings do not exclude the possibility that other classes of compounds may play a role in J2 host finding.

PPNs are highly damaging parasites with a tremendous impact on agriculture globally. *Meloidogyne* and *Globodera* spp. employ an extremely sophisticated and adaptive repertoire of effector molecules to subvert the plant host (reviewed by Mitchum et al,. 2013); the complexity and diversity of these organisms mean that crop Resistance (R) genes are either unavailable, or are insufficiently durable, to protect crops over the long term (reviewed by Davies and Elling, 2015). Reliance of synthetic chemical pesticides for their management can also have negative consequences to environmental and human health. Novel approaches, which are safe and effective, are required to control these parasites. Preventing the act of PPN host-finding and invasion represents an attractive intervention strategy for crop protection, and one that could be developed through manipulation of crop root exudate composition. This strategy has been effectively demonstrated against parasitic plants within the *Striga* genus (Mohemed et al., 2018; Xu et al., 2018; Gobena et al., 2017; Mohemed et al., 2016; Jamil et al., 2012), and could prevent secondary crop infection events. Our data demonstrate that crop plant gene expression can be modulated to alter exudate composition, as well as hatching and attraction of several important PPN species.

The GC-MS dataset reveals several compounds that are elevated in root exudates following ABC gene knockdown. When considering this relative to the behavioural response of PPN species to the exudates, we would hypothesise that elevated compounds function as repellents. Our behavioural data corroborate this for hexadecanoic acid, which is elevated following knockdown of *ABC-G33,* and pentadecane, which is elevated in exudates following knockdown of both *ABC-C6* and *ABC-G33.* Whilst our analysis of individual compounds has not been exhaustive, these observations do provide some confidence in our ability to predict exudate compound function using these approaches. It is clear however, that several of the compounds we selected for subsequent behavioural characterisation have no influence on PPN behaviour in these assays, and that complex interactions are likely involved. It should also be noted that root exudates will contain many additional compounds, which cannot be identified by GC-MS alone.

It is increasingly apparent that rhizosphere microbes are vital for plant health, and it has been shown that microbial chemotaxis is an important factor in the early colonisation of plant roots, and plant protection (Allard-Massicotte et al., 2016). We assessed the impact that root exudate compositional changes had on the positive chemotaxis of beneficial (*B. subtilis*) and pathogenic (*A. tumefaciens*) rhizosphere microbes to the experimental exudates. Our results indicate that *A. tumefaciens* was attracted to root exudates irrespective of the compositional changes following knockdown of either *ABC-C6* or *ABC-G33.* However, *B. subtilis* was significantly less attracted to root exudates following knockdown of *ABC-G33.* These data indicate that approaches to rhizosphere engineering, through the manipulation of root exudate composition, will need to assess a wide range of relevant organismal interactions. In this study, we used the domesticated *B. subtilis* strain 168 as an experimental model for *B. subtilis* chemotaxis. *B. subtilis* 168 does not form biofilms, unlike the undomesticated parental strain *B. subtilis* NCIB 3610 (Kesel et al., 2016; Zeigler et al., 2008). We found that *B. subtilis* NIB 3610 would form biofilms within our chemotaxis assay timeframe, making it difficult to enumerate individual cells that were attracted between experimental treatments. However, there may be additional implications for plant-microbe interactions, particularly in terms of biofilm and *vir* gene induction.

Our study relies upon a new approach to VIGS inoculation, which uses topical application of *A. tumefaciens* to seedling leaves, in a suspension containing the non-ionic surfactant and wetting agent, Silwet-L77. Unlike blunt syringe inoculation, this approach does not trigger growth stunting effects in the treated plant and promotes longer lasting gene knockdown. In the context of this study, using a transient gene silencing approach, such as VIGS has several conceptual benefits relative to other approaches that constitutively inhibit target gene expression. Transient knockdown can limit downstream secondary impacts in plants relative to constitutively inhibited genes by CRISPR-Cas9 or transgenic RNAi approaches, which could lead to false positive phenotypes. This phenomenon could develop as a function of biochemical knock-on effects that manifest phenotypically, but do not relate exclusively to target gene function, as previously suggested (Badri et al., 2008). Conceptually, we should have higher confidence in phenotypes recorded at earlier time points following loss of function analyses, which VIGS can facilitate. VIGS may also minimise genetic compensation, which occurs through transcriptional rescue of aberrant phenotypes, by counter-balanced expression of related genes. This can generate false negative phenotypes and is especially prominent in large gene families approaches (Rossi et al., 2015). Rossi et al. (2015) indicate that genetic compensation can be avoided or reduced using transient knockdown. VIGS typically results in a lower level of target gene knockdown than does transgenic dsRNA production and RNAi *in planta* (Albert et al., 2006), which may also reduce genetic compensation processes. VIGS is extremely cost-effective relative to other functional genomics approaches, in terms of both reagents and personnel time. It also represents the highest throughput reverse genetics tool available across a number of crop plants, and can be used to rapidly probe parasite interactions (Dubreuil et al., 2009). VIGS can also be used to silence multiple genes simultaneously (Orzaez et al., 2009).

Collectively, the data generated in this study support efforts to manipulate crop plant genes and promote beneficial rhizosphere interactions. This could occur through breeding, or biotechnology, and could support the sustainable intensification of global agriculture, through the rational and targeted exploitation of crop metabolic potential.

## Materials and Methods

### Virus Induced Gene Silencing

*Solanum lycopersicum* (cv. Moneymaker) seedlings were sterilised by rinsing in 1% sodium hypochlorite (from a diluted commercial bleach) for no more than 1 minute. Seeds were then rinsed in sterile ddH_2_O three times for no more than 2 minutes per wash. ddH_2_O was removed and seeds were sown on 0.5x MS agar plates (half strength Murashige and Skoog (MS) basal salt mixture, 2 mM Morpholinoethanesulfonic acid (MES), 1.5% agar (w/v), pH 5.7). Seeds were stratified at 4°C for 48 h in darkness before transfer to 23°C with 16 h of white light (140 – 160 μE.m^−1^.s^−1^)/8 h. Seedlings were transferred to John Innes number 2 compost upon cotyledon emergence.

The tobacco rattle virus (TRV) VIGS vector, pTRV2, was modified to contain a 200 bp fragment of the tomato *PDS* gene (Solyc03g123760; pTRV2-*PDS*). We used a co-silencing system as previously described (Orzaez et al., 2009; Stratmann & Hind, 2011), by generating pTRV2-*PDS-ABC-C6* and pTRV2-*PDS-ABC-G33* plasmids containing contiguous 200 bp fragments of the *PDS* gene sequence, followed by 200 bp of the gene of interest (either *ABC-C6* [Solyc08g006880] or *ABC-G33* [Solyc01g101070]). *Agrobacterium tumefaciens* strain GV3101 was used for VIGS throughout. *A. tumefaciens* cultures were transformed by electroporation, and stored as single use glycerol stocks. Briefly, 30 ng plasmid was added to 50 μl thawed electro-competent *A. tumefaciens* cells on ice, and then gently mixed by pipette. A single pulse of 2.2 kV was delivered to bacteria in a pre-chilled 1 mm gap cuvette. Cells were suspended in 1 ml LB broth, and incubated at 28°C for two hours at 200 rpm, before plating 50 μl on LB agar plates with 50 μg/ml kanamycin and 50 μg/ml gentamycin. Colonies were screened for successful transformation by colony PCR using universal pTRV backbone primers (see Table 1). Plants were inoculated with pTRV1/pTRV2 on the third day after transfer, between 2 – 4pm. From this point, plants were covered with a foil-lined propagator (to maintain humidity) for approximately 18 h at 18°C; the lower temperature is necessary to promote VIGS efficacy.

**Table 1.**
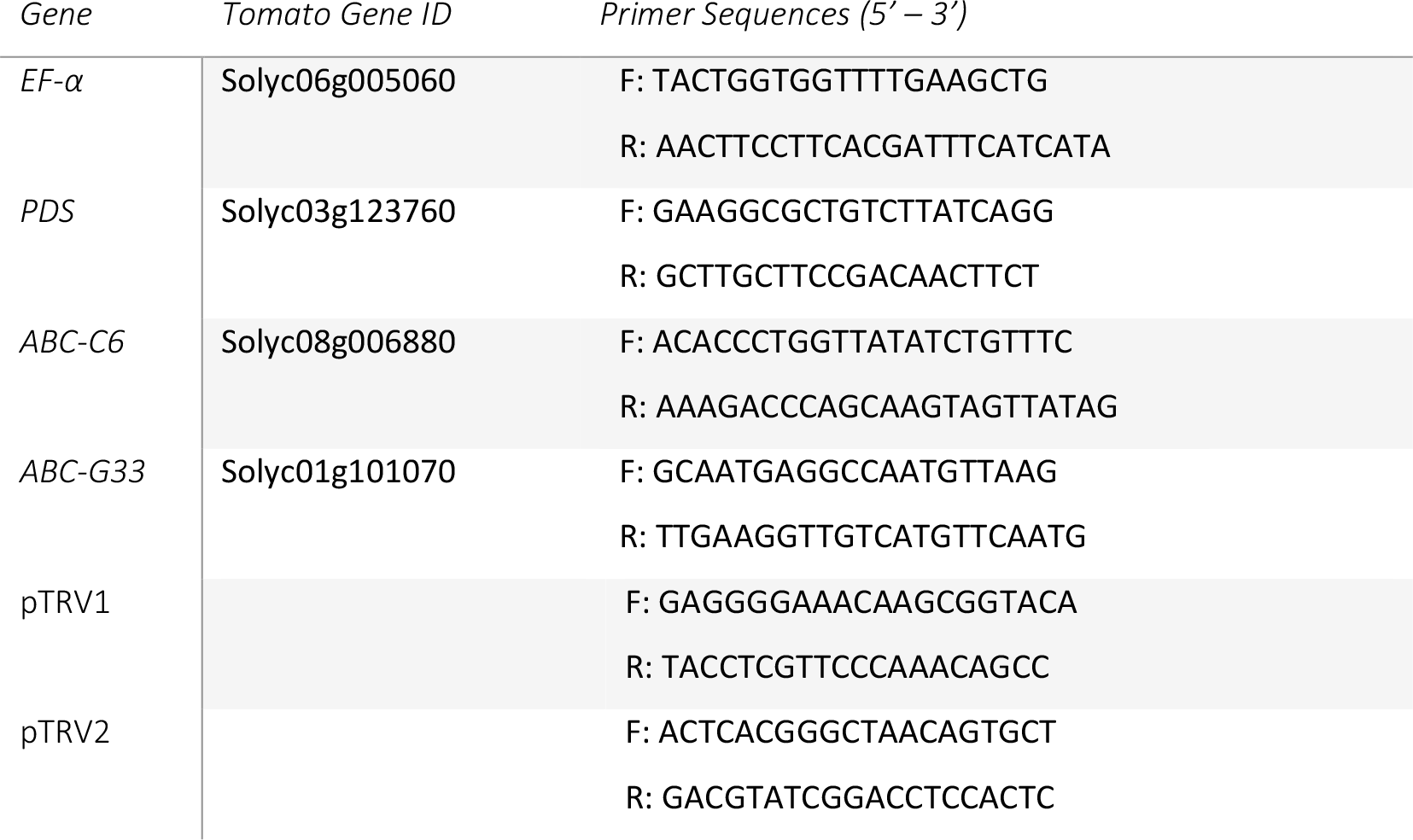
qRT-PCR primer sequences

### Preparation of *A. tumefaciens* cultures for tomato inoculation

For each construct, a single use glycerol stock was thawed and inoculated into 5 ml of LB broth containing 50 μg/ml kanamycin and 50 μg/ml gentamycin. pTRV1 was divided into two cultures (2 x 5 ml). Cultures were then incubated in darkness for 24-48 h at 28°C, with orbital agitation at 180 rpm, to an OD_600_ of between 0.75 – 1. *A. tumefaciens* cultures were then diluted to a total volume of 50 ml containing 50 μg/ml kanamycin and 50 μg/ml gentamycin, 10 mM MES, and 20 μM acetosyringone. Cultures were then incubated for 24 h at 28°C, with orbital agitation at 180 rpm, after which they were normalised to an OD_600_ of 1 in infiltration buffer (200 μM acetosyringone, 10 mM MES, 10 mM MgCl_2_, pH 5.7). Cultures were covered with foil and incubated for 3 h at room temperature. Immediately before topical application, a pTRV1 culture was mixed in a 1:1 v/v ratio with pTRV2, pTRV2-*PDS*, pTRV2-*PDS-ABC-C6*, or pTRV2-*PDS-ABC-G33*, as appropriate. Silwet-L77 was then added to a final concentration of 0.02%. No Silwet-L77 was added for the blunt syringe leaf infiltration method.

### Leaf infiltration and topical application methods

Prior to inoculation, the plants were watered to approximately 0.7 Field Capacity (FC). Leaf infiltration by blunt syringe was conducted as previously described, with a total volume of 0.1 ml injected per plant (Liu et al., 2002; Senthil-Kumar & Mysore, 2014). For topical application, 0.02% Silwett was added to the cultures immediately prior to application. An autoclaved paintbrush (size 10) was used to apply the culture across both abaxial and adaxial surfaces of the cotyledons, as well as the hypocotyl. Five strokes were administered to each seedling with a freshly inoculated paintbrush. Inoculated plants were maintained at 18°C, and covered with a foil-lined propagator for approximately 18 h. The propagator lid was then removed, and routine maintenance resumed at 18°C with a 16 h light, 8 h dark cycle, as before.

### Plant phenotype analysis

On the day of *A. tumefaciens* inoculation, and each subsequent day for six days, photographs were taken of five randomly selected plants in each treatment group to track growth of cotyledons. During this period cotyledon growth was found to be linear, after which growth begins to plateau and true leaves emerge. Thus the rate of cotyledon growth can be expressed as a function of y = mx + c. The rate of growth of each leaf was used in one-way ANOVA and Tukey’s HSD post-hoc tests to compare treatments. Plants were checked daily for leaf bleaching phenotypes, indicative of PDS co-silencing. The number of plants with unambiguous bleaching phenotypes were recorded daily and expressed as a percentage of the total number of plants inoculated.

### qRT-PCR analysis of gene transcript knockdown

Plants with bleaching phenotypes were removed from their pots and washed thoroughly under running water to remove soil. Three plants comprised one biological replicate in each treatment group at the times indicated. The plant tissue was wrapped in tin foil and flash frozen. Tissue was then crushed with a pestle and mortar into a fine powder. Frozen tissue was transferred to a 1.5 ml plastic microcentrifuge tube and total RNA was extracted using the Simply RNA Purification kit and Maxwell 16 extraction robot, following the manufacturer’s instructions (Promega). 10 μg total RNA was treated with Turbo DNase according to the manufacturer’s instructions (Ambion). 1 μg of purified RNA was subsequently reverse transcribed into cDNA using the High Capacity RNA-to-cDNA Kit as per manufacturer’s instructions (Applied Biosystems). A reverse transcription reagent (RTr) control, and a reverse transcription minus the reverse transcripase (RT-) control for a randomly selected sample per batch, were included.

cDNA, and controls, were diluted 1/4 using nuclease free water. 2.5 μl template was used for each qRT-PCR reaction in a total of 12 μl with 666 nM of each primer and 1x SensiFAST SYBR No-ROX mix as per manufacturer’s instructions (BIOLINE). Technical PCR reactions for each sample were performed in triplicate for each target using a Rotorgene Q thermal cycler with the following regime: [95°C x 10 min, 45 x (95°C x 20s, 60°C x 20s, 72°C x 25s) 72°C x 10 min]. PCR efficiencies of each amplicon, and the corresponding cT value, were calculated using the Rotorgene Q software. Relative quantification of each target amplicon was obtained by an augmented comparative Ct method (Pfaffl, 2001), relative to the reference gene EF-α (Solyc06g005060). Ratio-changes in transcript abundance were calculated relative to pTRV2-PDS treated plants. Data were analysed by one-way ANOVA and Tukey’s HSD post hoc test. Oligonucleotide sequences are listed in Table 1.

### Exudate collection

Plants were removed from their pots as described above, and cleared of soil by rinsing under running water. Five plants comprised one biological replicate. Plants were bunched together and their roots placed inside a 50 ml screw-top centrifuge tube (Corning), containing 10 ml ddH20. Plants were maintained under a standard 16 h light, 8 h dark regime at 18°C for the duration of exudate collection. After 24 h, at around 1 pm, plants were removed from the tubes and the exudate was passed through a 0.22 μM filter to remove root border cells and residual soil. Soil control samples (approximately 1 g soil in 10 ml 0.22 μM filtered ddH20) were processed in the same way. For chemosensory and hatching experiments, exudate was stored in centrifuge tubes at 4°C in darkness until use. For GC-MS metabolomics, exudates were stored at −80°C immediately after filtering.

### Non-targeted exudate metabolite profiling by GC-MS

Root exudates were freeze dried in batches and stored at −80°C until all samples had been processed (n = 40; 10 biological replicates for each treatment group). Samples were extracted with GC-grade dichloromethane (1 mL) (Sigma–Aldrich, St. Louis, MO, USA), vortexed for 10 s, sonicated for 5 min, and centrifuged at 14,000 rpm for 5 min. The organic phase was dried over anhydrous Na_2_SO_4_, concentrated to 50 μL under a gentle stream of N2 and then analysed (1.0 μL) by GC-MS on a 7890A gas chromatograph linked to a 5975 C mass selective detector (Agilent Technologies, Inc., Santa Clara, CA, USA). The GC was fitted with a HP5 MS low bleed capillary column (30 m × 0.25 mm i.d., 0.25 μm) (J&W, Folsom, CA, USA). Helium at a flow rate of 1.25 ml min-1 served as the carrier gas. The oven temperature was programmed from 35 to 285°C with the initial temperature maintained for 5 min then 10°C min-1 to 280°C, held at this temperature for 20.4 min. The mass selective detector was maintained at ion source temperature of 230°C and a quadrupole temperature of 180°C. Electron impact (EI) mass spectra were obtained at the acceleration energy of 70 eV. Fragment ions were analyzed over 40–550 m/z mass range in the full scan mode. The filament delay time was set at 3.3 min. A HP Z220 SFF intel xeon workstation equipped with ChemStation B.02.02. acquisition software was used. The mass spectrum was generated for each peak using Chemstation integrator set as follows: initial threshold = 5, initial peak width = 0.1, initial area reject = 1 and shoulder detection = on. The compounds were identified by comparison of mass spectrometric data and retention times with those of authentic standards and reference spectra published by library–MS databases: National Institute of Standards and Technology (NIST) 05, 08, and 11.

### Nematode culture and maintenance

*Meloidogyne* nematode species were maintained on *S. lycopersicum* (cv. Moneymaker). Four-week old plants were infected with 1,000 J2s. Eggs were extracted eight weeks post infection by first gently rinsing the root tissue free of soil and placing the tissue in a 500 ml Duran bottle. Tissue was then rinsed in 50% bleach for no more than one minute, and then in water, pouring the liquid through nested 180 μM, 150 μM, and 38 μM sieves arranged from largest pore size to smallest on the bottom. For maximum recovery, plant material was then crushed by hand and rinsed in water through the sieves. Eggs collected in the bottom sieve were pelleted and re-suspended in saturated sucrose. 1 ml ddH_2_0 was added to the solution, and eggs were collected by flotation in the water layer by centrifugation at 2000 rpm for 2 minutes. Eggs were collected and placed in a fine mesh filter hatchery (pore size 38 μm) in 5 ml spring water (autoclaved and sterilised by 0.22 μm filter). 50 μL antibiotic / antimycotic solution (Sigma) was added and eggs were incubated at 23°C. Freshly hatched J2s were collected every second or third day in hydrophobically lined microcentrifuge tubes for use in chemosensory and infection assays within 48 h of hatch.

*Globodera pallida* (pathotype PA2/3) cysts were reared on field grown potatoes (cv. Desiree). Two 10 week-old tomato (cv. Moneymaker) plants were grown under greenhouse conditions. 1 L ddH20 was poured into the soil and root stock of each pot, and collected after passing through the pot. Additional ddH20 was poured through the pots until the total collection volume reached 1 L. The collected exudate solution was passed through a 0.22 μM filter and stored in glass jars at 4°C until use. For chemosensory assays and infection assays, *G. pallida* cysts were hatched in 1:1 (v/v) tomato exudate diluted in ddH20 and collected at two to three-day intervals in hydrophobically lined microcentrifuge tubes.

### Soil infection assays

Tomato plants exhibiting the co-silenced leaf bleaching phenotype were selected at around 21 days post inoculation, and then challenged with 250 J2s. All treatments were blinded from this stage. Six weeks after J2 inoculation, galls and cysts could be identified. These were counted for each experimentally infected plant.

### PPN chemosensory assays

Chemosensory assays were conducted as before (Warnock et al., 2016; 2017) by making a solid agar base (3 ml of 1.5% (w/v) agar) in a 60 mm Petri dish. 3 ml of smooth 0.5% (w/v) agar slurry was then poured on top of the solid base. Slurry was prepared by continuous mixing of liquid 0.5% agar until fully cooled. Approximately 150 J2s were then added to the centre of the arena under a microscope. Plates were covered and left in darkness at room temperature (20°C) overnight on a vibration free bench (for approx. 16 h). The numbers of nematodes observed in the different regions of the arena were then counted under a light microscope. The chemosensory index of each assay was calculated as before (Warnock et al., 2016; 2017).

### PPN egg hatching assays

Tomato root exudate (collected as described above) was diluted in a 1:1 ratio with ddH2O. 500 μl of diluted exudate was dispensed into wells of a 24-well culture plate. To each exudate sample, between 15 and 20 *G. pallida* cysts were added. Four biological replicates were prepared for each treatment group. The spaces between wells were half filled with ddH20 and the plates wrapped in parafilm to reduce evaporation and changes of volume throughout the experiment. Plates were incubated at 17°C for 21 days. Each well was checked daily for signs of nematode emergence. Following the first day of emergence, J2s were counted every 48 h. After approximately 3 weeks, the remaining unhatched eggs / J2s were counted in order to obtain a cumulative percentage hatch rate for each exudate sample.

The number of emerging J2s at each time point was converted to a cumulative percentage, which was plotted against time as previously described (Campbell & Madden, 1990).

2000 freshly extracted *M. incognita* or *M. javanica* eggs were incubated in 500 μl of experimental plant root exudates. The ratio of unhatched to hatched J2s was then recorded every second or third day. These ratio values were converted into percentages and plotted relative to time, as previously described. For each batch of eggs, triplicate counts of unhatched and hatched J2s were made. Four experimental replicates were used to calculate means.

### Microbe chemotaxis assays

Microbial chemotaxis assays were conducted broadly as in Allard-Massicotte et al. (2016). *B. subtilis* (168) and *A. tumefaciens* (AGL-1) were inoculated onto LB agar plates and spread to single colonies. One day-old colonies were inoculated into 3 ml LB broth and orbitally rotated at 180 rpm, overnight (28°C for *A. tumefaciens* and 37°C for *B. subtilis*). Cultures were pelleted at 8,000 rpm for 10 min, and the cells were washed in 1.5 ml chemotaxis buffer (10 mM Potassium Phosphate Buffer, pH 7.0), 0.1 mM EDTA, 0.05% glycerol, 5 mM Sodium-d,l-Lactate, 0.14 mM CaCl_2_, 0.3mM (NH_4_)_2_SO_4_). Cells were collected by centrifugation and subsequently re-suspended in fresh chemotaxis buffer to an OD_600_ of 0.002. 200 μl of cell suspension was added to each well of a 96-well plate; ten replicates for each experimental group.

1 μl microcapillary tubes (Sigma-Alrdich) were filled with either: (i) experimental root exudates (*ABC-C6, ABC-G33,* or *PDS*), (ii) positive (1 mM malic acid), or negative (ddH_2_O) controls. Loaded microcapillary tubes were placed into the cell suspension wells of the 96-well plate for 1 h, and maintained at 23°C. During this time, planktonic *B. subtilis* or *A. tumefaciens* cells could migrate towards, and into, the microcapillary tube. Following the 1 h assay timecourse, the capillary tubes were removed. Excess cell suspension was removed from the outside of each capillary tube by rinsing briefly with ddH_2_O. The 1 μl content of each capillary tube was ejected into 99 μl of chemotaxis buffer by positive pressure. 20 μl of each solution was spread onto a 1.5% LB agar plate. LB plates were sealed with parafilm, and incubated at 28°C for *A. tumefaciens*, or 37°C for *B. subtilis* for 48 h. Colony forming units were counted for each replicate plate.

## Acknowledgements

VIGS plasmids pTRV1 and pTRV2 were kindly provided by Prof. Ian Graham, University of York. *A. tumefaciens* AGL-1 was provided by Dr Vladimir Nekrasov, Rothamsted Research. *A. tumefaciens* strain GV3101 was provided by Dr Penny Hirsch, Rothamsted Research.

## References

Adebesin F, Widhalm JR, Boachon B, Lefèvre F, Pierman B, Lynch JH, Alam I, Junqueir B, Benke R, Ray S, Porter JA, Yanagisawa M, Wetzstein HY, Morgan JA, Boutry M, Schuurink RC, Dudareva N (2017). Emission of volatile organic compounds from petunia flowers is facilitated by an ABC transporter. Science, 356(6345): 1386–1388. https://doi.org/10.1126/science.aan0826

Allard-Massicotte R, Tessier L, Lecuyer F, Lakshmanan V, Lucier JF, Garneau D, Caudwell L, Vlamakis H, Bais HP, Beauregard PB (2016). Bacillus subtilis early colonization of Arabidopsis thaliana roots involves multiple chemotaxis receptors. MBio 7(6): e01664–16. https://doi.org/10.1128/mBio.01664-16

Ahkami AH, White RA, Handakumbura PP, Jansson C (2017). Rhizosphere engineering: enhancing sustainable plant ecosystem productivity. Rhizosphere, 3(2): 233–243. https://doi.org/10.1016/j.rhisph.2017.04.012

Badri DV, Loyola-Vargas VM, Broeckling CD, De-la-Pena C, Jasinski M, Santelia D, Martinoia E, Sumner LW, Banta LM, Stermitz F, Vivanco JM (2008). Altered profile of secondary metabolites in the root exudates of Arabidopsis ATP-Binding Cassette (ABC) transporter mutants. Plant Physiol. 146(2): 762–771. https://doi.org/10.1104/pp.107.109587

Badri DV, Quintana N, El Kassis EG, Kim HK, Choi YH Sugiyama A, Verpoorte R, Martinoia E, Manter DK, Vivanco JM (2009). An ABC transporter mutation alters root exudation of phytochemicals that provoke an overhaul of natural soil microbiota. Plant Physiol. 151(4):2006–2007. https://doi.org/10.1104/pp.109.147462

Berendsen RL, Pieterse CMJ and Bakker PAHM (2012). The rhizosphere microbiome and plant health. Trends in Plant Sci 17(8): 478–486. https://doi.org/10.1016/j.tplants.2012.04.001

Bertin C, Yang XH, Weston LA (2003). The role of root exudates and allelochemicals in the rhizosphere. Plant Soil 256, 67–83. https://doi.org/10.1023/A:1026290508166

Bowsher AW, Ali R, Harding SA, Tsai CJ, Donovan LA (2016). Evolutionary divergences in root exudate composition among ecologically-contrasting Helianthus species. PLoS One 11(1): e1048280: https://doi.org/10.1371/journal.pone.0148280

Bulgarelli D, Garrido-Oter R, Munch PC, Weiman A, Droge J, Pan Y, McHardy AC, Schulze-Lefert P (2015). Structure and function of the bacterial root microbiota in wild and domesticated barley. Cell Host Microbe. 17(3): 392–403. https://doi.org/10.1016/j.chom.2015.01.011

Byrne JT, Maher NJ, Jones PW (2001). Comparative responses of Globodera rostochiensis and G. pallida to hatching chemicals. J Nematol 33(4): 195–202. DOI not available.

Campbell CL, Madden LV (1990). Introduction to Plant Disease Epidemiology. John Wiley & Sons, New York. ISBN: 0471832367.

Chaparro JM, Badri DV, Bakker MG, Sugiyama A, Manter DK, Vivanco JM (2013). Root exudation of phytochemicals in Arabidopsis follows specific patterns that are developmentally programmed and correlate with soil microbial functions. PLoS ONE 8(8): https://doi.org/10.1371/journal.pone.0055731

Cieslinksi G, Van Rees CJ, Szmigielska AM, Huang PM (1997). Low molecular weight organic acids released from roots of durum wheat and flax into sterile nutrient solutions. J Plant Nutr 20: 753–764. https://doi.org/10.1080/01904169709365291

Davies LJ, Elling AA (2015). Resistance genes against plant-parasitic nematodes: a durable control strategy? Nematology 17(3): https://doi.org/10.1163/15685411-0000287.

De Andrade LRM, Ikeda M, do Amaral LIV, Ishizuka J (2011). Organic acid metabolism and root excretion of malate in wheat cultivars under aluminium stress. Plant Physiol Biochem 49(1): 55–60. https://doi.org/10.1016/j.plaphy.2010.09.023

Dessaux Y, Grandclement C, Faure D (2016). Enginerring the rhizosphere. Trends in Plant Sci 21(3): 266–278. https://doi.org/10.1016/j.tplants.2016.01.002

de Weert S, Vermeiren H, Mulders IH, Kuiper I, Hendrickx N, Bloemberg GV, Vanderleyden J, De Mot R, Lugtenberg BJ (2002). Flagella-driven chemotaxis towards exudate components is an important trait for tomato root colonization by Pseudomonas fluorescens. Mol Plant Microbe Interact 15(11): 1173–80. https://doi.org/10.1094/MPMI.2002.15.11.1173

Dubreuil G, Magliano M, Dubrana MP, Lozano J, Lecomte P, Favery B, Abad P, Rosso MN (2009). Tobacco rattle virus mediates gene silencing in a plant parasitic root-knot nematode. J Exp Bot 60(14): 4041–4050. https://doi.org/10.1093/jxb/erp237

Duarte A, Maleita C, Abrantes I, Curtis R (2015). Tomato root exudates induce transcriptional changes of Meloidogyne hispanica genes. Phytopathol Mediterr 54: 1, 104−108. https://doi.org/10.14601/Phytopathol_Mediterr-14595

Fletcher JS, Hegde RS (1995). Release of phenols by perennial plant roots and their potential importance in bioremediation. Chemosphere 31(4): 3009–3016. https://doi.org/10.1016/0045-6535(95)00161-Z

Fudali SL, Wang C, Williamson VM (2013). Ethylene signalilng pathway modulates attractiveness of host roots to the root-knot nematode Meloidogyne hapla. Mol Plant Microbe Interact 26(1): 75–86. https://doi.org/10.1094/MPMI-0512.0107-R

Giles CD, Brown LK, Adu MO, Mezeli MM, Sandral GA, Simpson RJ, Wendler E, Shand CA, Menezes-Blackburn D, Darch T, Stutter MI, Lumsdon DG, Zhang H, Blackwell MSA, Wearing C, Cooper P, Havgarth PM, George TS (2017). Response-based selection of barley cultivars and legume species for complementarity: Root morphology and exudation in relation to nutrient source. Plant Sci 255: 12–28. https://doi.org/10.1016/j.plantsci.2016.11.002

Gobena D, Shimels M, Rich PJ, Ruyter-Spira C, Bouwmeester H, Kanuganti S, Mengiste T, Ejeta G (2017). Mutation in sorghum LOW GERMINATION STIMULANT 1 alters strigolactones and causes Striga resistance. PNAS 114(17): 4471–4476. https://doi.org/10.1073/pnas.1618965114

Hiltpold I, Baroni M, Toepfer S, Kuhlmann U, Turlings TCJ (2010). Selection of entomopathogenic nematodes for enhanced responsiveness to a volatile root signal helps to control a major root pest. J Exp Biol 213: 2417–2423. https://doi.org/10.1242/jeb.041301

Hubbard CJ, Li B, McMinn R, Brock MT, Maignien L, Ewers BE, Kliebenstein D, Weinig C (2018). The effect of rhizosphere microbes outweighs host plant genetics in reducing insect herbivory. Mol Ecol in press https://doi.org/10.1111/mec.14989

Hu Y, You J, Li C, Williamson VM, Wang C (2017). Ethylene response pathway modulates attractiveness of plant roots to soybean cyst nematode Heterodera glycines. Sci Rep 7: 41282. https://doi.org/10.1038/srep41282

Hwang J-U, Song W-Y, Hong D, Ko D, Yamaoka Y Jang S, Yim S, Lee E, Khare D, Kim K, Palmgren M, Yoon HS, Martinoia E, Lee Y (2016). Plant ABC transporters enable many unique aspects of a terrestrial plant’s lifestyle. Mol Plant 9(3): 338–355. https://doi.org/10.1016/j.molp.2016.02.003

Iannucci A, Fragasso M, Beleggia R, Nigro F, Papa R (2017). Evolution of the crop rhizosphere: impact of domestication on root exudates in tetraploid wheat (Triticum turgium L.). Front Plant Sci 8: 2124. https://doi.org/10.3389/fpls.2017.02124

Jamil M, Charnikhova T, Houshyani B, van Ast A, Bouwmeester HJ (2012). Genetic variation in strigolactone production and tillering in rice and its effect on Striga hermonthica infection. Planta 235(3): 473–484. https://doi.org/10.1007/s00425.1520-y

Jones JT, Haegeman A, Danchin EGJ, Gaur HS, Helder J, Jones MGK, Kikuchi T, Manzanilla-Lopez R, Palomares-Ruis JE, Wesemael WML, Perry RN (2013). Top 10 plant-parasitic nematodes in molecular plant pathology. Mol Plant Pathol 14(9): 946–961. https://doi.org/10.1111/mpp.12057

Jones DL, Nguyen C, Finlay RD (2009). Carbon flow in the rhizosphere: carbon trading at the soil-root interface. Plant Soil 321, 5–33. https://doi.org/10.1007/s11104-009-9925-0

Kesel S, Grumbein S, Gumperlein I, Tallawi M, Marel A-K, Lieleg O, Opitz M (2016). Direct comparison of physical properties of Bacillus subtilis NCIB 3610 and B-1 biofilms. Appl Environ Microbiol 82: 2424–2432. https://doi.org/10.1128/AEM.03957-15.

Kihika R, Murungi LK, Coyne D, Ng’ang’a M, Hassanali A, Teal PEA, Torto B (2017). Parasitic nematode Meloidogyne incognita interactions with different Capsicum annum cultivars reveal the chemical constituents modulating root herbivory. Sci Rep 7: 2903. https://doi.org/10.1038/s41598-017-02379-8

Kim B, Song GC, Ryu CM (2016). Root exudation by aphid leaf infestation recruits root-associated Paenibacillus spp. to lead plant insect susceptibility. J Microbiol Biotechnol 26(3): 549–57. https://doi.org/10.4014/jmb.1511.11058

Kirwa HK, Murungi LK, Beck JJ, Torto B (2018). Elicitation of Differential Responses in the Root-Knot Nematode Meloidogyne incognita to Tomato Root Exudate Cytokinin, Flavonoids, and Alkaloids. J Agric Food Chem 66(43): 11291–11300. https://doi.org/10.1021/acs.jafc.8b05101

Lane TS, Rempe CS, Davitt J, Staton ME, Peng Y, Soltis DE, Melkonian M, Deyholos M, Leebens-Mack JH, Chase M, Rothfels CJ, Stevenson D, Graham SW, Yu J, Liu T, Pires JC, Edger PP, Zhang Y, Xie Y, Zhu Y, Carpenter E, Wong GK-S, Stewart CN (2016). Diversity of ABC transporter genes across the plant kingdom and their potential utility in biotechnology. BMC Biotechnol 16(1): 47. https://doi.org/10.1186/s12896-016-0277-6

Li B, Li YY, Zhang FF, Li CJ, Li XX, Lambers H, Li L (2016). Root exudates drive interspecific facilitation by enhancing nodulation and N2 fixation. PNAS 113(23): 6496–501. https://doi.org/10.1073/pnas.1523580113

Liu G, Sanches-Fernandez R, Li Z-S, Rea PA (2001). Enhanced multispecificity of Arabidopsis vacuolar multidrug resistance-associated protein-type ATP-binding cassette transporter, AtMRP2. J Biol Chem 276: 8648–8656. https://doi.org/10.1074/jbc.M009690200

Lu T, Ke M, Lavoie M, Jin Y, Fan X, Zhang Z, Fu Z, Sun L, Gillings M, Penuelas J, Qian H, Zhu YG (2018). Rhizosphere microorganisms can influence the timing of plant flowering. Microbiome 6(1): 231. https://doi.org/10.1186/s40168-018-0615-0

Marschner H (1995). Mineral nutrition of higher plants (2nd Edition). Academic Press. ISBN: 9780124735439.

Martinoia E, Meyer S, De Angeli A, Nagy R (2012). Vacuolar transporters in their physiological context. Annu Rev Plant Biol 63: 183–213. https://doi.org/10.1146/annurev-arplant-042811-105608.

Meyer RS, Duval AE, Jensen HR (2012). Patterns and processes in crop domestication: an historical review and quantitative analysis of 203 global food crops. New Phytol 196(1): 29–48. https://doi.org/10.1111/j.14698137.2012.04253.x

Micallef SA, Shiaris MP, Colon-Carmona A (2009). Influence of Arabidopsis thaliana accessions on rhizobacterial communities and natural variation in root exudates. J Exp Bot 60: 1729–1742. https://doi.org/10.1093/jxb/erp053

Mitchum MG, Hussey RS, Baum TJ, Wang X, Elling AA, Wuben M, Davis EL (2013). Nematode effector proteins: an emerging paradigm of parasitism. New Phytol 199: 879–894. https://doi.org/10.1111/nph.12323

Mohemed N, Charnikhova T, Fradin EF, Rienstra J, Babiker AGT Bouwmeester HJ (2018). Genetic variation in Sorghum bicolor strigolactones and their role in resistance against Striga hermonthica. J Exp Bot https://doi.org/10.1093/jxb/ery041

Mohemed N, Charnikhova T, Bakker EJ, van Ast A, Babiker AG, Bouwmeester HJ (2016). Evaluation of field resistance to Striga hermonthica (Del.) Benth. In Sorghum bicolor (L.) Moench. The relationship with strigolactones. Pest Manag Sci 71(11): 2082–2090. https://doi.org/10.1002/ps.4426

Monchgesang S, Strehmel N, Trutschel D, Westphal L, Neumann S, Scheel D (2016). Plant to plant variability in root metabolite profiles of 19 Arabidopsis thaliana accessions is substance-class-dependent. Int J Mol Sci 17(9): E1565. https://doi.org/10.3390/ijms17091565-

Monchgesang S, Strehmel N, Schmidt S, Westphal L, Taruttis F, Muller E, Herklotz S, Neumann S, Scheel D (2017). Natural variation of root exudates in Arabidopsis thaliana-linking metabolomic and genomic data. Sci Rep 6, Article number: 29033. https://doi.org/10.1038/srep29033

Morris R, Wilson L, Warnock ND, Carrizo D, Cox D, Sturrock M, Maule AG, Dalzell JJ (2017). A neuropeptide modulates sensory perception in the entomopathogenic nematode Steinernema carpocapsae. PLoS Pathog 13(3): e1006185. https://doi.org/10.1371/journal.ppat.1006185

Murungi LK, Kirwa H, Coyne D, Teal PEA, Beck JJ, Torto B (2018). Identification of key root volatiles signaling preference of tomato over spinach by the root knot nematode Meloidogyne incognita. J Agric Food Chem 66: 7328–7336. https://doi.org/10.1021/acs.jafc.8b03257

Nicol JM, Stirling GR, Turner SJ, Coyne DL, de Nijs L, Hockland S, Maafi ZT (2011). Current nematode threats to world agriculture. Genomics and Molecular Genetics of Plant-Nematode Interactions (Jones JT, Gheysen G, Fenoll C., eds). Heidelberg: Springer pp. 21–44. https://doi.org/10.1007/97894.007-0434-3-2

Ofori PA, Mizuno A, Suzuki M, Martinoia E, Reuscher S, Aoki K, Shibata D, Otagaki S, Matsumoto S, Shiratake K (2018). Genome-wide analysis of ATP binding cassette (ABC) transporters in tomato. PLoS ONE 13(7): e0200854. https://doi.org/10.1371/journal.pone.0200854

Palomares-Rius JE, Hedley P, Cock PJA, Morris JA, Jones JT, Blok VC (2016). Gene expression changes in diapause or quiescent potato cyst nematode, Globodera pallida, eggs after hydration or exposure to tomato root diffusate. PeerJ 4: e1654. https://doi.org/10.7717/peerj.1654

Rossi A, Kontarakis Z, Gerri C, Nolte H, Holper S, Kruger M, Stainier DYR (2015). Genetic compensation induced by deleterious mutations but not gene knockdowns. Nature 524: 230–233. https://doi.org/10.1038/nature14580

Runyon JB, Mescher MC, De Moraes CM (2006). Volatile chemical cues guide host location and host selection by parasitic plants. Science 313: 1964–1967. https://doi.org/10.1126/science.1131371

Sasse J, Martinoia E, Northen T (2018). Feed your friends: do plant exudates shape the root microbiome? Trends in Plant Sci 23(1): 25–41. https://doi.org/10.1016/j.tplants.2017.09.003

Schroeder JI, Delhaize E, Frommer WB, Guerinot ML, Harrison MJ, Herrera-Estrella L, Horie T, Kochian LV, Munns R Nishizawa NK, Tsay Y-F, Sanders D (2013). Using membrane transporters to improve crops for sustainable food production. Nature 497: 60–66. https://doi.org/doi:10.1038/nature11909

Teillet A, Dybal K, Kerry BR, Miller AJ, Curtis RHC, Hedden P (2013). Transcriptional changes of the root-knot nematode Meloidogyne incognita in Response to Arabidopsis thaliana root signals. PLoS ONE 8(4): e61259. https://doi.org/10.1371/journal.pone.0061259

Warnock ND, Wilson L, Canet-Perez JV, Fleming T, Fleming CC, Maule AG, Dalzell JJ (2016). Exogenous RNA interference exposes contrasting roles for sugar exudation in host-finding by plant pathogens. Int J Parasitol 46(8): 473–7. https://doi.org/10.1016/j.ijpara.2016.02.005

Warnock ND, Wilson L, Patten C, Fleming CC, Maule AG, Dalzell JJ (2017). Nematode neuropeptides as transgenic nematicides. PLoS Pathog 13(2): e1006237. https://doi.org/10.1371/journal.ppat.1006237

Waters BM, Amundsen K, Graef G (2018). Gene expression profiling of iron deficiency chlorosis sensitive and tolerant soybean indicates key roles in phenylpropanoids under alkalinity stress. Front Plant Sci 9: 10. https://doi.org/10.3389/fpls.2018.00010

Winzer T, Gazada V, He Z, Kaminski F, Kern M, Larson TR, Meade F, Teodor R, Vaistij FE, Walker C, Bowser TA, Graham IA (2012). A Papaver somniferum 10-gene cluster for synthesis of the anticancer alkaloid noscapine. Science 336: 1704–1708. https://doi.org/10.1126/science.1220757

Xu Y, Miyakawa T, Nosaki S, Nakamura A, Lyu Y, Nakamura H, Ohto U, Ishida H, Shimizu T, Asami T, Tanokura M (2018). Structural analysis of HTL and D14 proteins reveals the basis for ligand selectivity in Striga. Nat Commun 9(1): 3947. https://doi.org/10.1038/s41467-018-06452-2

Yang G, Zhou B, Zhang X, Zhang Z, Wu Y, Zhang Y, et al. Effects of tomato root exudates on Meloidogyne incognita (2016). PLoS ONE 11(4): e0154675. https://doi.org/10.1371/journal.pone.0154675

Yazaki K (2006). ABC transporters involved in the transport of plant secondary metabolites. FEBS Lett 580(4): 1183–91. https://doi.org/10.1016/j.febslet.2005.12.009

Yuan J, Zhang N, Huang Q, Raza W, Li R, Vivanco JM, Shen Q (2015). Organic acids from root exudates of banana help root colonization of PGPR strain Bacillus amyloliquefaciens NJN-6. Sci Rep 5: 13438. https://doi.org/10.1038/srep13438

Zasada IA, Peetz A, Wade N, Navarre RA, Ingham RE (2016). Host status of different potato (Solanum tuberosum) varieties and hatching in root exudates of Globodera ellingtonae. J Nematol 45(3): 195–201. DOI not available. PMID: 24115784

Zeigler DR, Pragai Z, Rodriguez S, Chevreux B, Muffler A, Albert T, Bai R, Wyss M, Perkins JB (2008). The origins of 168, W23, and other Bacillus subtilis legacy strains. J Bacteriol 190(21): 6983–6995. https://doi.org/10.1128/JB.00722-08.

Zwetsloot MJ, Kessler A, Bauerle TL (2018). Phenolic root exudate and tissue compounds vary widely among temperate forest tree species and have contrasting effects on soil microbial respiration. New Phytol 218(2): 530–541. https://doi.org/10.1111/nph.15041

